# Leveraging Experimental Evolution to Extract Predictive Collateral Drug Response Signatures in Ewings Sarcoma

**DOI:** 10.1101/2025.09.19.677399

**Authors:** Kristi Lin-Rahardja, Ethan Canavan, Jacob G Scott

## Abstract

Therapeutic options for patients with relapsed or refractory Ewings sarcoma (EWS) remain limited. Collateral sensitivity, where resistance to one drug confers sensitivity to another, could be leveraged to optimize chemotherapy for EWS. Gene expression signatures that predict collateral sensitivity states can be used to guide treatment selection in an evolution-informed manner. We experimentally evolved resistance to first-line EWS chemotherapy in cell lines. Throughout, we measured collateral responses across a panel of anticancer drugs and quantified transcriptomic changes. Collateral drug responses varied across replicate evolutionary trajectories, but convergent states of collateral sensitivity emerged across different replicates at different times. By associating these convergent phenotypes with gene expression patterns, we derived a library of predictive signatures for numerous drugs. These signatures accurately distinguished states of collaterally sensitivity from states of collateral resistance within our dataset and were further validated in an independent EWS cell line. Our findings demonstrate that gene expression signatures can predict collateral sensitivity in EWS, providing a foundation for personalized therapeutic strategies. This approach also provides a generalizable workflow for developing predictive biomarkers to guide chemotherapy selection in patients with rare diseases that lack reliable second-line chemotherapy regimens.

## Introduction

Ewings sarcoma (EWS) is a rare bone malignancy primarily observed in children and adolescents.^1,2^ A quarter of patients with localized disease and 70% of patients with metastatic disease ultimately relapse.^1,3–6^ The 5-year survival rate for relapsed patients is less than 15%.^3,4,7^ In addition to surgery and radiation, first-line treatment includes alternating combinations of vincristine, doxorubicin, and cyclophosphamide (VDC) with ifosfamide and etoposide (IE). Individuals requiring second-line chemotherapy typically receive temozolomide and cyclophosphamide or irinotecan, temozolomide, and vincristine. Like the first-line chemotherapy regimen, the second-line regimen combines alkylating agents with topoisomerase inhibitors. The reliance of both regimens on overlapping mechanisms of action makes cross-resistance difficult to avoid, but also presents a clear opportunity to vastly improve how we approach treatment for advanced EWS patients.

Predictive models for drug response would be highly beneficial for optimizing chemotherapy regimens in individual patients, especially for those with advanced EWS who have few effective chemotherapy options. One straightforward, easily translatable approach for predicting drug response is the use of gene signatures, or a set of genes whose expression is associated with increased sensitivity toward a particular drug. To better formulate a personalized chemotherapy regimen for patients with treatment-resistant EWS, we can extract signatures that specifically predict collateral sensitivity, a phenomenon in which resistance to one drug increases sensitivity to another. Utilizing this concept would be informative for guiding treatment strategies by informing clinicians which agents may be most effective against a resistant tumor after relapse.^8–10^ The concept of using collateral sensitivity to guide treatment has been explored for bacterial infections as well as cancer, and predictive gene signatures would be an attractive option for applying this strategy to a clinical oncology setting.^9,11–13^

Although EWS tumors are genetically homogeneous and have a low mutation rate, intratumoral transcriptomic heterogeneity is extensive.^2,14,15^ This heterogeneity can make it difficult to identify robust markers of drug-sensitive tumor states, but exploiting convergent evolution can help overcome this difficulty and yield more reliable predictors.^16,17^ Convergent evolution is the phenomenon in which independent populations evolve the same phenotype. In the context of predictive signature extraction, we can exploit this principle by identifying patterns of gene expression that emerge across different tumors or tumor subpopulations that all exhibit sensitivity against a drug. This approach has been demonstrated previously to extract a pan-cancer cisplatin sensitivity signature from two large datasets, the Genomics of Drug Sensitivity in Cancer (GDSC) and The Cancer Genome Atlas (TCGA), and has been validated with other clinical datasets.^16^ Unfortunately, data on drug efficacy against a wide variety of agents in EWS patients is extremely limited, especially for the subset of these patients with recurrent tumors who desperately need an advancement in treatment strategy. To generate the large amount of data needed to extract gene signatures predictive of convergent states of collateral sensitivity in treatment-resistant EWS, we performed experimental evolution.

We evolved resistance against first-line chemotherapy in multiple independent EWS cell line replicates, simultaneously quantifying collateral drug responses to a broad panel of 48 agents while capturing transcriptomic changes across evolutionary time. This approach allowed us to identify convergent drug-sensitive states despite the stochasticity of collateral drug responses, and to extract predictive biomarkers associated with these states. Ultimately, we generated 28 predictive gene signatures of collateral sensitivity and validated their utility in an independent EWS cell line. Beyond the immediate findings, the dataset we generated represents a rare, large-scale resource for modeling evolution-informed drug response in EWS and the workflow we outline is a generalizable method for other rare diseases lacking reliable second-line chemotherapy options.

## Results

### Resistance to first-line chemotherapy evolves to varying degrees in EWS cell lines

To evolve resistance against the first-line chemotherapy used against EWS, we first split a population of an EWS cell line, CHLA9, into ten *evolutionary replicates* to proliferate and evolve independently over time. Seven replicates (numbered evolutionary replicates/ER 1-7) were treated to drive chemoresistance, while the remaining three replicates (ER 8-10) were treated with the relevant concentrations of dimethyl sulfoxide (DMSO) and sodium thiosulfate as vehicle controls (**Fig.1A**). The seven experimental replicates were treated with alternating combinations of VDC (vincristine, doxorubicin, and cyclophosphamide) and EC (etoposide and cyclophosphamide). In clinical practice, patients with EWS receive etoposide and ifosfamide, not EC. However, ifosfamide cannot be metabolically activated *in vitro*. As such, we used cyclophosphamide in its place, which has the same mechanism of action and can be activated *in vitro* with sodium thiosulfate. The ratio between the drugs within each combination is based on the maximum achievable concentration in plasma for the individual agents.^18^ These ratios remained constant for all dose-response assays involving these two combinations. To drive resistance in the experimental replicates, we treated the cell lines with the *EC*_70_s of VDC and EC. We maintained a constant high-dose treatment throughout the experiment to mimic clinical administration of chemotherapy at the maximum tolerated dose. The replicates were treated for ten cycles, starting with VDC and alternating between EC at every subsequent time point.

**Figure 1.**
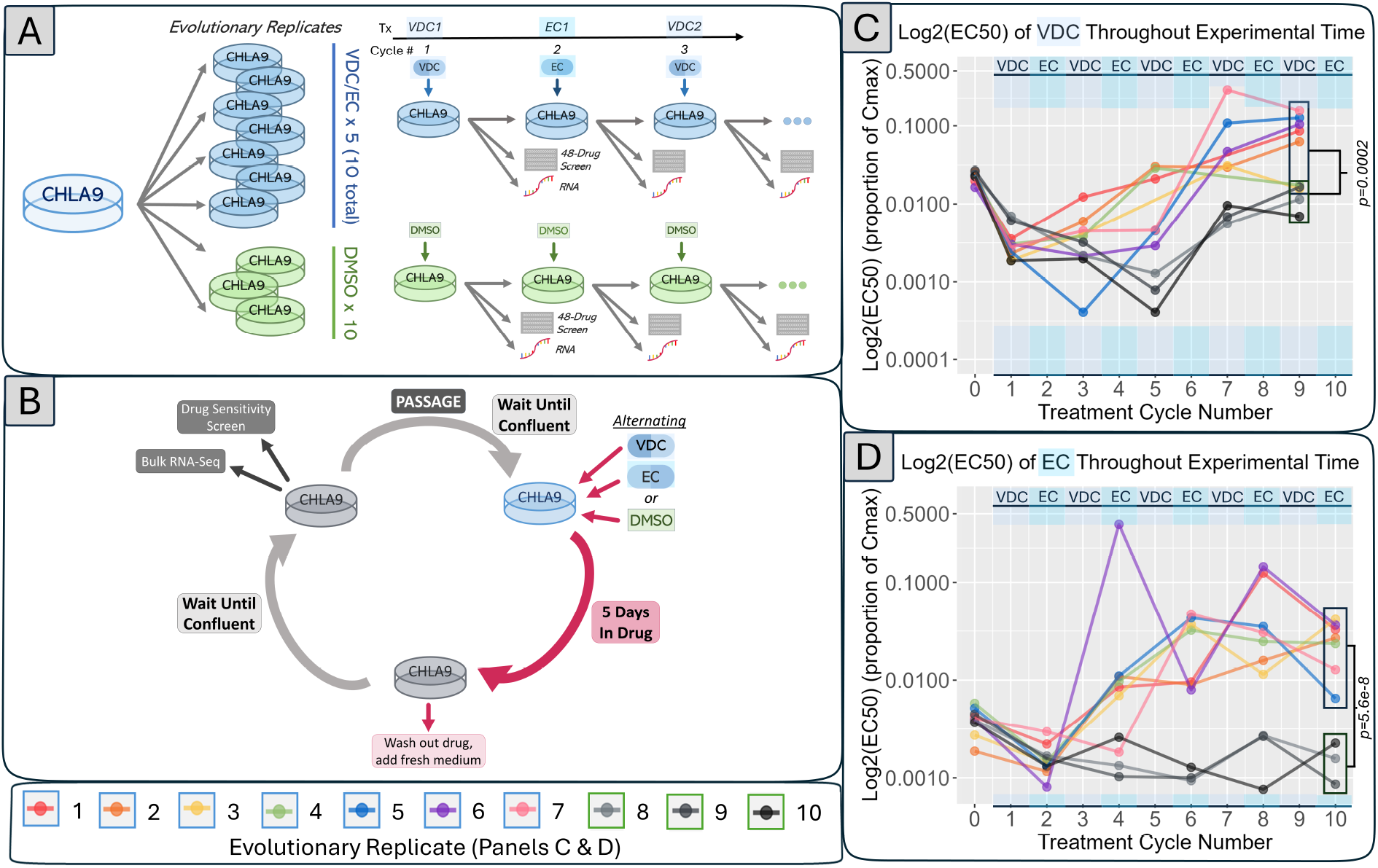
Experimental design for evolving drug resistance. **A. Overview of experimental method**. One population of CHLA9 was split into ten evolutionary replicates. Seven replicates (ER 1-7) were treated with alternating doses of VDC and EC for a total of ten exposures. The remaining three replicates (ER 8-10) served as a vehicle control for ten passages. After each treatment, each replicate was sampled for various assays and storage. **B. Detailed schematic of each treatment cycle**. Each cycle within this experiment proceeded as pictured. Samples were grown to confluency, exposed to drug for five days, allowed to regrow without drug to generate enough cells for testing, then harvested for various measurements and passaged for the subsequent cycle. **C and D. Resistance evolved to first-line chemotherapy over time**. For both panels, experimental time proceeds along the x-axis, and the treatment being administered at each time point is noted along the top of each plot. The log_2_(*EC*_50_) against VDC (panel **C**) or EC (panel **D**) is shown on the y-axis. The concentration unit of the *EC*_50_ is described as the proportion of the maximum concentration used in the assay for simplicity, since different concentrations are used for each drug (see *Materials & Methods* for exact concentrations). Experimental replicates are colored according to the legend, and control replicates are colored in grayscale. A Welch’s *t*-test between experimental and control replicates at the end of the experiment shows statistically significant differences in *EC*_50_s between the two groups for both drug combinations.

Each treatment cycle included a drug-free growth stage, five days of exposure to one of the two drug combinations, and then a second drug-free growth stage. Once the sample was confluent again, the replicate was harvested for drug response screening, RNAseq, freezing, and passaging to seed the next treatment cycle (**Fig.1B**).

Throughout the ten cycles, the experimental replicates became more resistant to both VDC and EC while the control replicates maintained a similar response for the entire duration of the experiment (**Fig. 1C and D**). The control replicates, while mostly stable, did become slightly more sensitive over time. Generally, the experimental replicates evolved a higher degree of resistance against EC than VDC. This result could be due to more drugs being included in VDC than EC. The increased complexity of the VDC selection pressure may have been more difficult for the cells to adapt to, resulting in a lesser degree of resistance compared to EC. For both combinations, different evolutionary replicates achieved varying degrees of resistance and, in some cases, showed fluctuations in the strength of resistance over time. These fluctuations could be attributed to the selection pressure changing every cycle.

To investigate overall changes in gene expression between resistant and treatment-naive EWS, we performed differential expression analysis between treatment-naive (time point 0, the ancestor) and resistant replicates (time point 10). Gene set enrichment analysis on differentially expressed genes revealed differences in the expression patterns between evolutionary replicates (**Fig. S1**).^19^ Comparisons between treatment-naive and end-point resistant replicates were done independently, since we hypothesized that each evolutionary replicate could evolve resistance through different means even under the same conditions. In comparing replicate two to treatment-naive CHLA9, IL6/JAK/STAT3 signaling and genes involved in apical junctions were enriched in the resistant sample. In replicate five, genes targeted by ZNF618 were enriched. In replicate six, genes involved in myogenesis were upregulated, while genes involved in hepatic metabolism and xenobiotic response were downregulated. These differences in gene expression among experimental replicates support our hypothesis that tumor cells of the same origin under the same selection pressure can evolve resistance in numerous ways, and despite these differences, they share a convergent phenotype of resistance towards VDC and EC. Future investigation of mutation status across the replicates would provide further insight into this matter. However, the primary objective of this study is not to elucidate mechanisms of VDC-EC resistance in EWS, but to identify transcriptomic biomarkers that clinicians can use to improve the treatment of patients with resistant tumors.

### Collateral drug response varies across replicates and time

All evolutionary replicates were tested against 48 other agents and the selection panel (VDC and EC) at the end of every treatment cycle. The drugs in the collateral response screen have all exhibited antitumor activity and employ various mechanisms of action (see **Supplemental File 1** for all agents, their mechanisms, and other details). This screen included 15 common chemotherapies used across multiple cancer types, including irinotecan, temozolomide, and topotecan, which are commonly used as second-line treatments for EWS.^20^ 13 targeted therapies are also included. In addition, 19 experimental cancer drugs were included. 7 of these experimental agents were developed specifically to treat cancer, including SP-2509, a KDM1A/LSD1 inhibitor designed to treat Ewings sarcoma.^21,22^ The remaining 12 experimental agents included drugs that could be repurposed to treat cancer. We quantified collateral responses against the drugs for each replicate at each time point by calculating the fold-change in drug response between the evolved sample and treatment-naive CHLA9.

Over time, all replicates consistently became more resistant against certain agents, such as gemcitabine and floxuridine (**Fig. 2**). The opposite was true for some other drugs, such as FX-11 and ML-210. The collateral drug screen revealed some instances of repeatable collateral resistance across drugs with similar mechanisms to the first-line therapy, such as paclitaxel and docetaxel. A majority of samples evolved collateral resistance to irinotecan, topotecan, and temozolomide, the drugs often used as second-line agents against EWS. This result is not unexpected due to the similarity in the mechanisms of action of these drugs compared to first-line chemotherapy. Doxorubicin and etoposide (first-line agents) are topoisomerase II inhibitors, and irinotecan and topotecan (second-line agents) are topoisomerase I inhibitors. Cyclophosphamide (a first-line agent) is an alkylating agent, as is temozolomide (a second-line agent).

**Figure 2.**
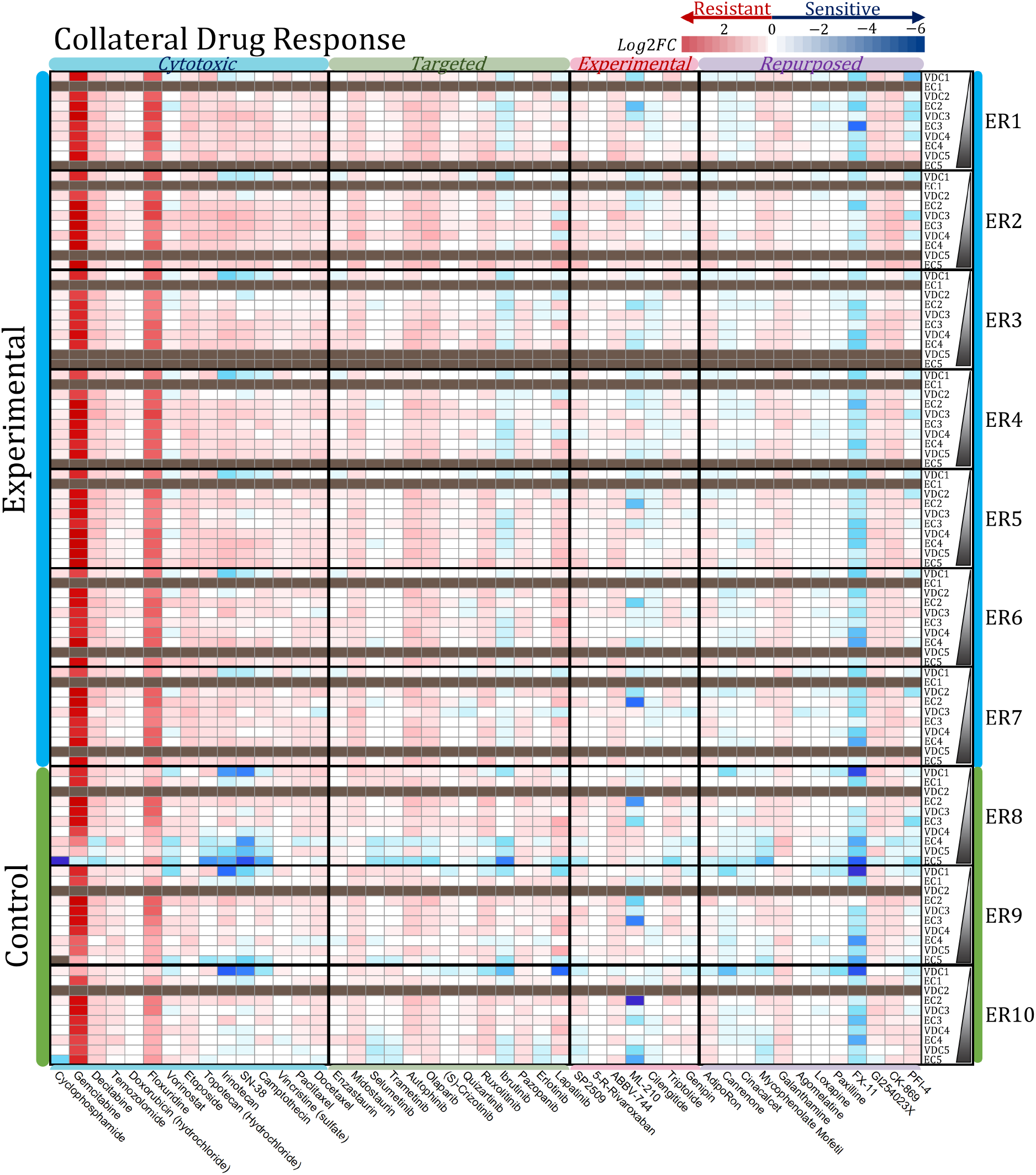
Collateral drug response over time. Drug response to each agent is shown in each column. Columns are grouped by overall drug category as indicated at the top of the plot. Rows are grouped by evolutionary replicate, with experimental time proceeding from top to bottom within each group. Log_2_ fold-change in response compared to treatment naive, or time point 0, is represented colorimetrically, where increased resistance toward an agent is red and increased sensitivity is blue. Brown cells indicate missing data due to contamination.

In several cases, similarly strong collateral drug responses evolved in both the experimental replicates and control replicates, most noticeably to gemcitabine, floxuridine, and FX-11. These results imply that these particular collateral responses may be driven by mechanisms independent of VDC/EC resistance. However, it should also be noted that the control replicates’ collateral resistance towards gemcitabine and floxuridine decreases toward the end of the experiment, but it is maintained in the experimental replicates. The collateral sensitivity to FX-11 in the control replicates is also somewhat stronger than in the experimental replicates. Thus, while the primary driving forces for these similar response profiles may not be directly related to the mechanisms driving VDC-EC resistance, those mechanisms could still be influencing these collateral responses to some degree.

As a whole, the majority of drugs tested in the collateral screen showed varying levels of efficacy across the experiment (**Fig. 3**). We expected this result because a change in one trait (i.e., collateral sensitivity or resistance against a drug) is independent of selection acting on a different trait (i.e., evolution under VDC/EC exposure).^23^ Repeatable instances of collateral sensitivity or resistance may suggest that these agents can reliably be used or should be avoided. Predictive gene signatures would be most useful in cases where there is a wider distribution of collateral response across tumors, and it is not obvious whether the disease would be sensitive or resistant at any point in time.

**Figure 3.**
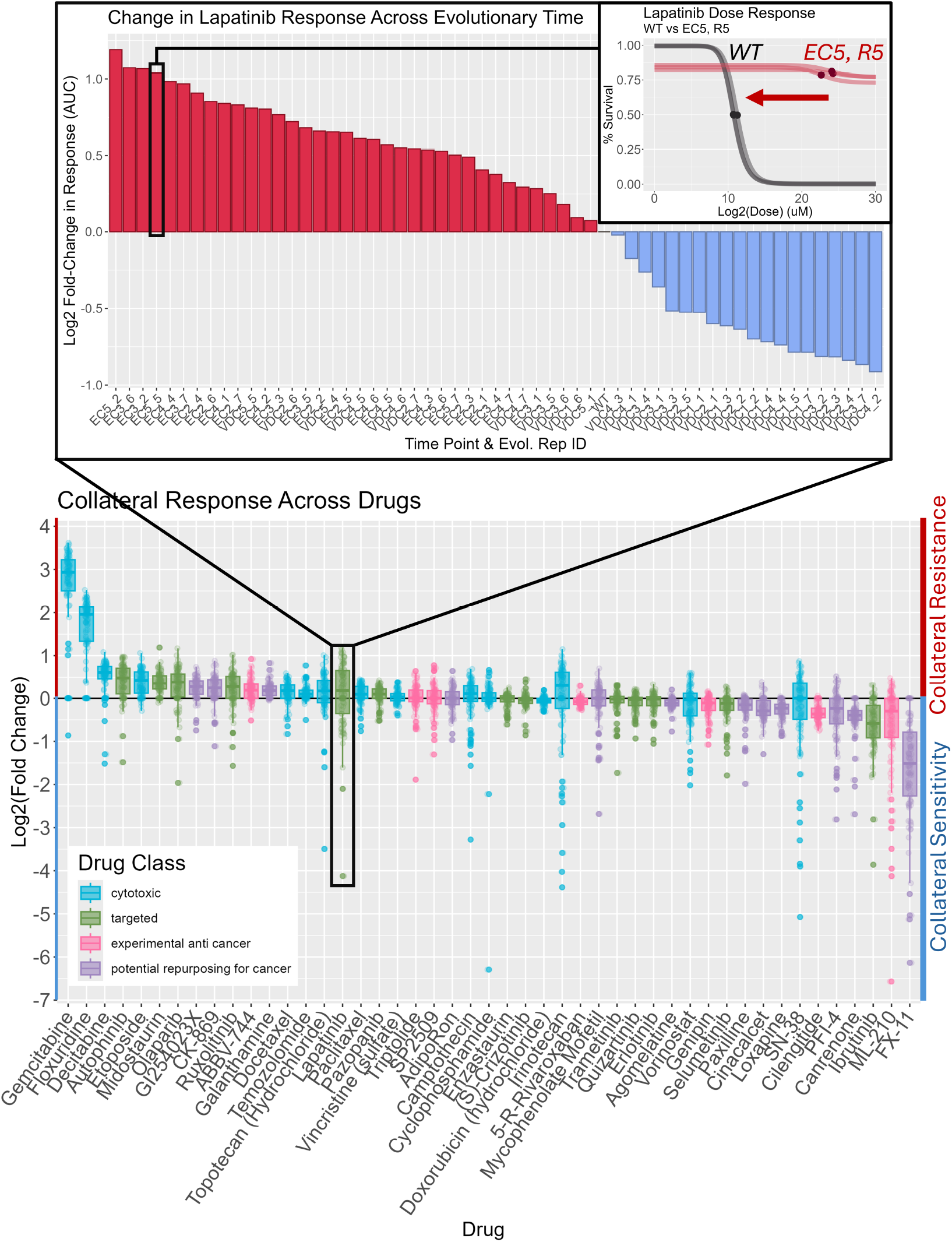
Collateral responses across experimental replicates and time. Change in collateral drug response is measured by comparing drug response between wild-type treatment-naive CHLA9 and the sample of interest, as shown in the top right. The top waterfall plot shows collateral drug response for one of the drugs in the screen, lapatinib, stratified by individual samples from most resistant to most sensitive, irrespective of experimental time. The lower plot depicts collateral responses across all 48 drugs, and each point within a box-whisker represents one experimental evolutionary replicate at one point in time. Drugs included in the collateral screen are listed along the x-axis by descending median log_2_(fold-change in AUC). Drugs toward the left (with positive log_2_ fold-changes) show samples with higher collateral resistance across all samples, and drugs toward the right (with negative log_2_ fold-changes) show samples with higher collateral sensitivity across all samples. Boxplots for each drug are colored by general drug category. **7/27**

### Comparing differential gene expression between sensitive and resistant samples to extract predictive signatures of collateral sensitivity

To extract predictive gene signatures for each drug in the collateral response screen, we compared gene expression between samples that showed the most sensitivity and the most resistance relative to the treatment-naive sample, regardless of replicate identity and experimental time. By selecting resistant and sensitive samples regardless of treatment history, we aimed to identify convergent states of drug sensitivity and find gene expression patterns similarly reflected across these disparate samples. To increase stringency and reduce the false discovery rate, we performed 5-fold repeated subsampling, where we first divided the pool of samples into five groups of equal size. Then, using four of five folds, we conducted differential expression analysis to compare the top and bottom 20% of samples ordered by drug response within the given subset of data.^24^ This process was repeated five times, with a different fold removed from the analysis each time. The overexpressed genes extracted by EBSeq in at least three of the five folds were retained in the final signature (**Fig. 4**). From this analysis, we extracted 35 predictive signatures, each associated with sensitivity to one of the agents in the collateral drug screen. The process was run for all 48 drugs, but 13 yielded no genes, leaving us with 35 gene signatures. We repeated this pipeline using only experimental replicates and also resulted in 35 gene signatures, although this set of signatures differed by six drugs compared to the initial set of signatures. Signatures unique to the first set (extracted from all replicates) included mycophenolate mofetil, ABBV-744, vincristine, ML-210, erlotinib, and GI254023X. Signatures unique to the second set (extracted from experimental replicates) included paclitaxel, cinacalcet, triptolide, quizartinib, midostaurin, and cyclophosphamide.

**Figure 4.**
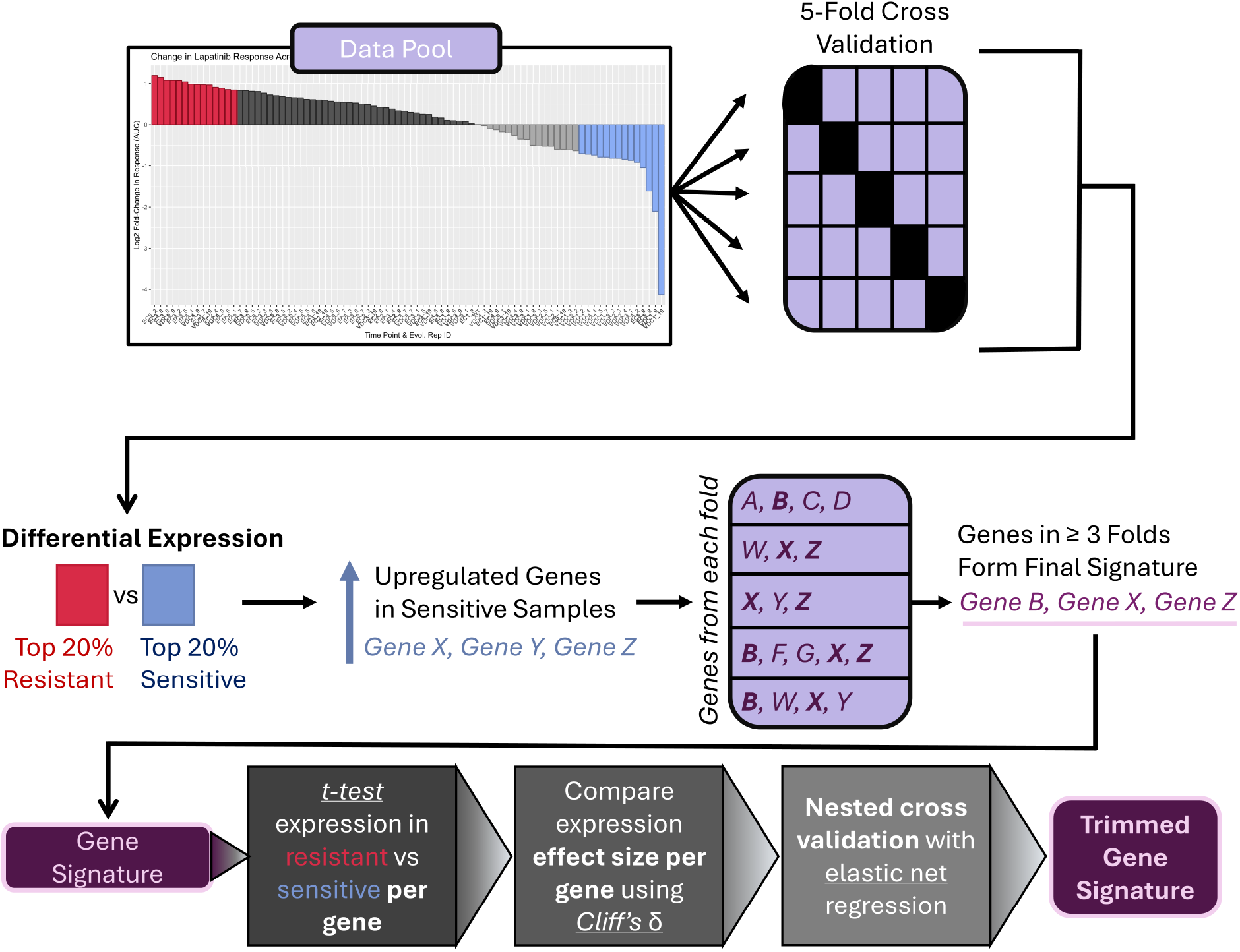
Signature extraction method. For each drug included in the collateral drug screen, we extracted a gene signature to predict collateral drug response. Predictive signatures were extracted by performing 5-fold repeated subsampling, where all sequenced samples were split evenly into five folds. Differential expression between the top 20% most resistant and sensitive samples was performed five times, with a different fold excluded from the analysis each time. Only the genes found to be upregulated in the sensitive samples were kept. The genes extracted in at least three of the five folds were retained to form the final signature. Signatures were extensively trimmed using various statistical approaches (see **Methods** for further details).

Several signatures included hundreds of genes, which would be impractical for future clinical translation. As such, all existing signatures went through extensive trimming using various statistical metrics. For each gene in a signature, we performed a *t*-test on the expression between the 20% most resistant and 20% most sensitive samples and kept only the genes with a statistically significant difference after Bonferroni correction. For the remaining genes, we calculated Cliff’s Delta between the same sample groups using a conservative threshold of |*δ* | > 0.8.^25^ Lastly, we fitted an elastic net regularized logistic regression with nested cross-validation (see **Materials & Methods** for further details).^26,27^ As a result of these three trimming methods, seven of the 35 signatures extracted from all replicates were removed entirely (decitabine, mycophenolate mofetil, gemcitabine, doxorubicin, topotecan, docetaxel, and GI254023X), leaving 28 trimmed signatures. For the signatures extracted from experimental replicates only, the doxorubicin and 5-R-rivaroxaban signatures were removed, resulting in 33 trimmed signatures. The number of genes in the signatures before and after trimming is shown in **Figs. S3** and**S4**.

### Within-sample testing of signature efficacy

To assess the performance of our signatures within our dataset, we fit numerous statistical models to predict survival using the expression of each gene in a signature as a predictor in the model (**Fig. 5A**). Four linear models and seven non-linear models were trained for each signature, using a randomly selected 50 out of 72 sequenced samples to perform leave-one-out cross-validation. The linear models were trained to predict the percentage of surviving cells at the highest concentration tested against the corresponding drug. The non-linear models were trained to predict whether the percentage of surviving cells at the highest measured dose was above or below the median survival across all samples. To quantify the accuracy of the models, we used the remaining 22 to test the models and compared the predicted outcome to the observed outcome. For linear models, we calculated Spearman’s rho between these values, and for the non-linear models, we calculated the area under the receiver operating curve (AUC).

**Figure 5.**
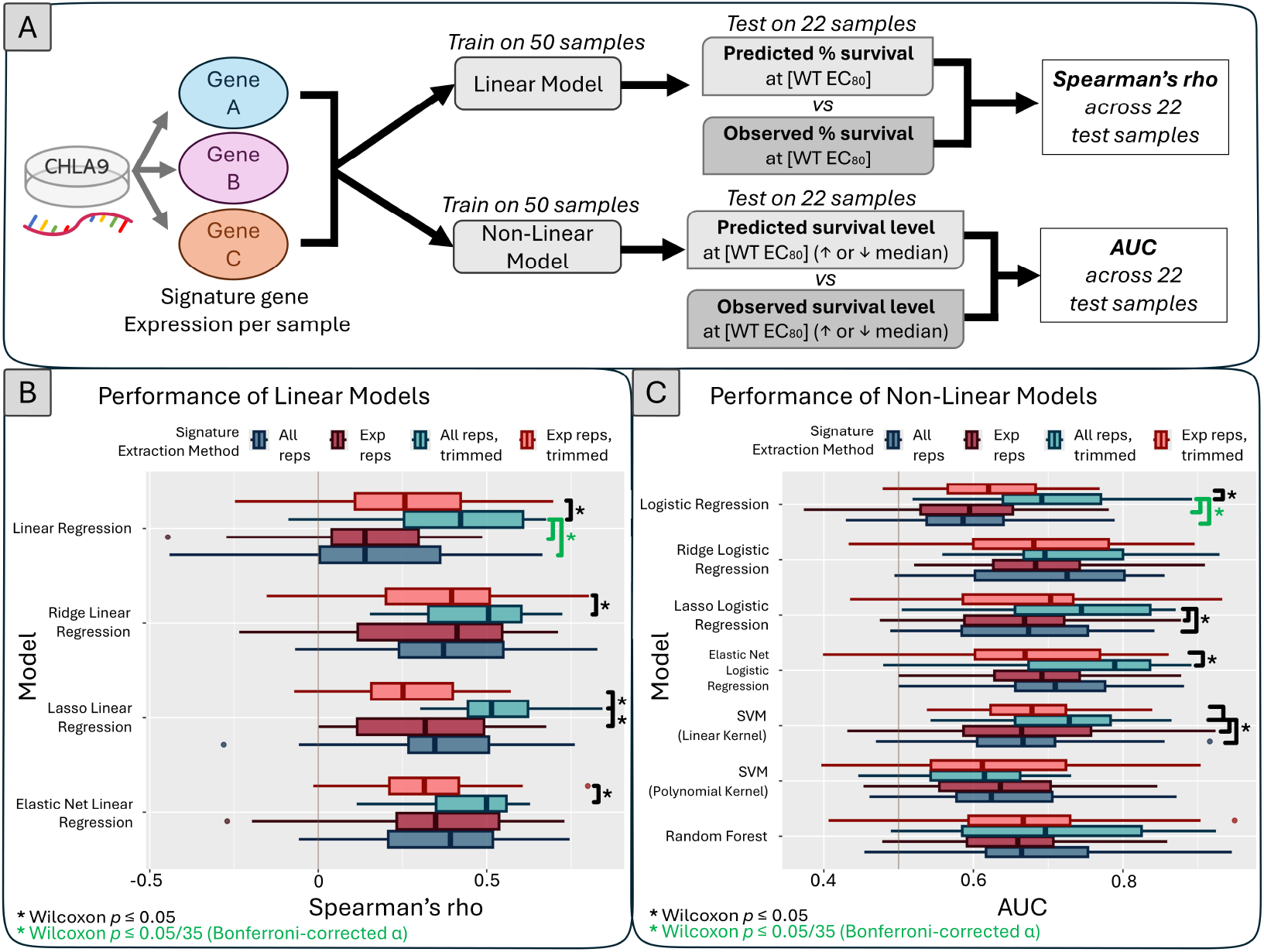
Signature genes are used as predictors in various statistical models to assess performance. A. Schematic of the modeling pipeline. We trained each model on 50 random samples from the 72 we had sequenced, using the expression of individual signature genes as predictors. The trained models are then tested on the remaining 22 samples that were not used for training. Linear models predict the percent of surviving cells at the wild-type cell line’s *EC*_80_. Non-linear models predict whether the percent survival at that dose is above or below the median survival across all samples. Spearman’s rho or area under the receiver operating curve (AUC) is then calculated by comparing predicted and observed survival. **B. Spearman’s rho is compared among signatures extracted from different methods**. Spearman’s rho is indicated along the x-axis, and the model used is shown on the y-axis. Each boxplot depicts the distribution of Spearman’s rho from all signatures. Different colors indicate different extraction methods. Statistical significance between pairs of boxplots is indicated by the black and green brackets. **C. AUC is compared among signatures extracted from different methods**. This panel is laid out in the same format as **panel B**, but depicts AUC along the x-axis.

Before analyzing the accuracy of individual signatures, we first compared how different extraction methods would influence overall performance. The different methods we used included extraction from all replicates (control and experimental) with or without trimming, and extraction from experimental replicates with or without trimming. Spearman’s rho and AUC across all signatures, grouped by extraction method, are shown in **Figs. 5B and C**. Across all but one model, signatures extracted from all replicates with trimming outperform signatures extracted from the other three attempted methods, suggesting that this method would be the most robust moving forward. Additionally, we compared how different extraction methods performed between signatures that included only up-regulated genes in sensitive samples or signatures that included both up- and down-regulated genes enriched in sensitive samples (**Figs. S5 and S6**). Across all models, we found no major performance differences between up-regulated signatures and up- and down-regulated signatures together, and none of these comparisons yielded statistical significance. Since the addition of the down-regulated gene signatures yielded no significant improvement to prediction accuracy and because gene up-regulation is more reliably detectable than down-regulation and tends to be more statistically robust, we chose to move forward with the up-regulated gene signatures only.^28,29^ From this point on, we will focus on testing the 28 trimmed signatures extracted from all replicates whose expression is up-regulated when samples are sensitive (see **Supplemental File 2** for lists of genes included in each of these signatures).

To more thoroughly assess the performance of individual signatures, we compared Spearman’s rho and AUC derived from our signatures to a null distribution. For each signature, we generated 200 random signatures of the same length and used these genes as predictors to perform the same pipeline illustrated in **Fig. 5A**. The metrics generated from this bootstrap formed the null distributions shown in **Figs. 6A** (example of one signature), **S7**, and **S8** (results for all signatures). To summarize these comparisons across all signatures and models, we computed an effect size (*z*-score) by standardizing our signatures’ performance (Spearman’s rho or AUC) relative to their corresponding null distribution. These *z*-scores are visualized as a heatmap in **Fig. 6B**.

**Figure 6.**
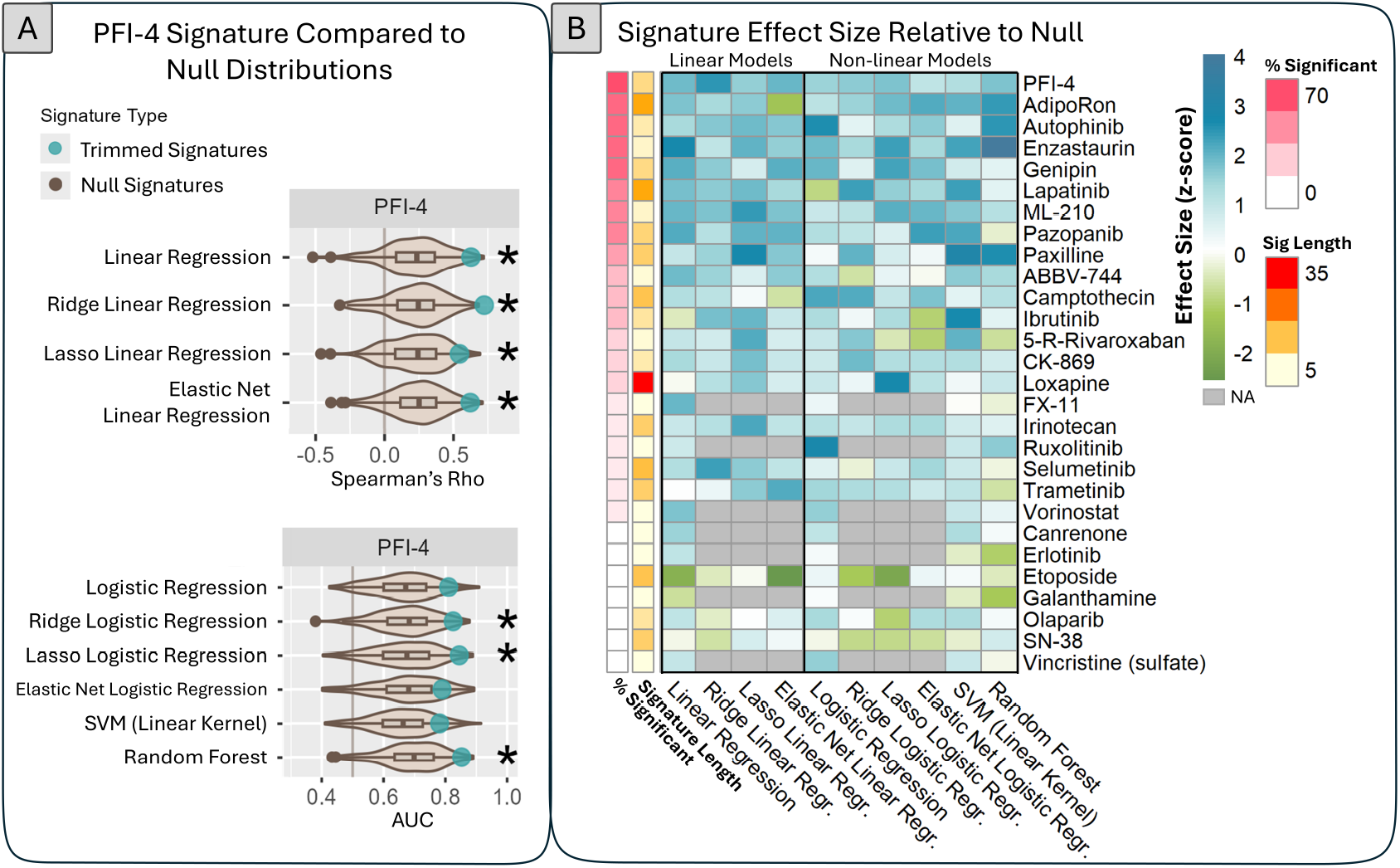
Comparison of signature performance to null distributions. **A. Example of null distributions generated for one signature, applied to all models**. Performance metrics of the PFI-4 signature are shown as blue points. The null distribution is shown in beige. Empirical p-values were calculated for each model, and p-values ≤ 0.05 are indicated with an asterisk. These comparisons for all other signatures are shown in **Figs. S7 and S8. B. Effect size for each signature and model, relative to the null distribution**. Effect size is calculated as a *z*-score of the signature’s Spearman’s rho or AUC relative to the corresponding null distribution. A *z*-score *>* 0 indicates that the signature yields more accurate predictions than the null distribution. A *z*-score *<* 0 indicates that the signature performed worse than the null distribution. Signatures are shown by row, and the various models are shown by column. Z-scores are indicated in blue (positive) and green (negative). Some penalized linear and logistic models were unable to be fitted with the random gene signatures, resulting in missing data indicated by gray cells in the heatmap. Additional annotations for the signatures are shown along the left of the primary heatmap. The leftmost (pink) column indicates the percentage of models for which the signature’s performance was significantly different from its null distribution. The adjacent (orange) column indicates how many genes are in the signature.

Across 10 models trained on our 28 signatures, we observed a positive *z*-score 85% of the time, indicating that the majority of our signatures could predict drug sensitivity more accurately than random genes. This result was not the case for the etoposide signature, as suggested by the consistently negative z-score and lack of statistical significance across models. This result was not surprising, however, given that etoposide was included in the first-line treatment for EWS, and nearly all samples were resistant against this drug to some degree. As such, there were likely not enough samples to properly identify genes that are up-regulated when a sample is sensitive to etoposide. There was no correlation between the length of a signature and its ability to make accurate predictions about drug response.

### External validation of signatures

A critical step to translate our signatures to a clinical setting will be external validation. In order to perform such an analysis, we must have gene expression data paired with drug response against the 28 drugs for which we have extracted signatures. Using data obtained from this study’s pilot experiment by Scarborough *et al*, we validated three of our signatures.^17^ In this pilot study, we evolved resistance against first-line chemotherapy in a different EWS cell line, A673, for ten cycles. The collateral drug response screen included twelve drugs, and we sequenced eighteen samples from varying time points throughout the experiment. Our collateral screen included eleven of these twelve drugs; we did not include dactinomycin in this experiment. Because we were unable to extract a signature for all the drugs we tested in our screen, we had only five applicable signatures for the A673 dataset. Two of these signatures (vorinostat and vincristine) included only one gene per signature, so they were excluded from this analysis. The three remaining signatures we were able to test included etoposide (12 genes), olaparib (6 genes), and SN-38 (10 genes).

To assess the predictive capability of these three signatures in A673, we compared signature scores between the 20% most resistant and 20% most sensitive samples against the drug in question. The signature score is represented by the median normalized expression of the genes in a given signature, a common summarization method for such biomarkers.^30^ Since the expression of our signatures should be increased when a sample is sensitive, we expected that the sensitive samples would have a higher signature score than the resistant samples. This expectation was the case for both CHLA9 and A673 (**Fig. 7A and B**). The differences in signature scores observed between response groups in CHLA9 were statistically significant for all three signatures. While this was not the case for A673, the differences between groups were distributed as expected, with sensitive samples scoring higher than resistant samples. Additionally, we compared the signature scores to the *EC*_50_s of all 18 A673 samples. Since signature scores should be higher with increased sensitivity and vice versa, we expected the scores to decrease as *EC*_50_ increased, and this was true for all three signatures (**Fig. 7C**).

**Figure 7.**
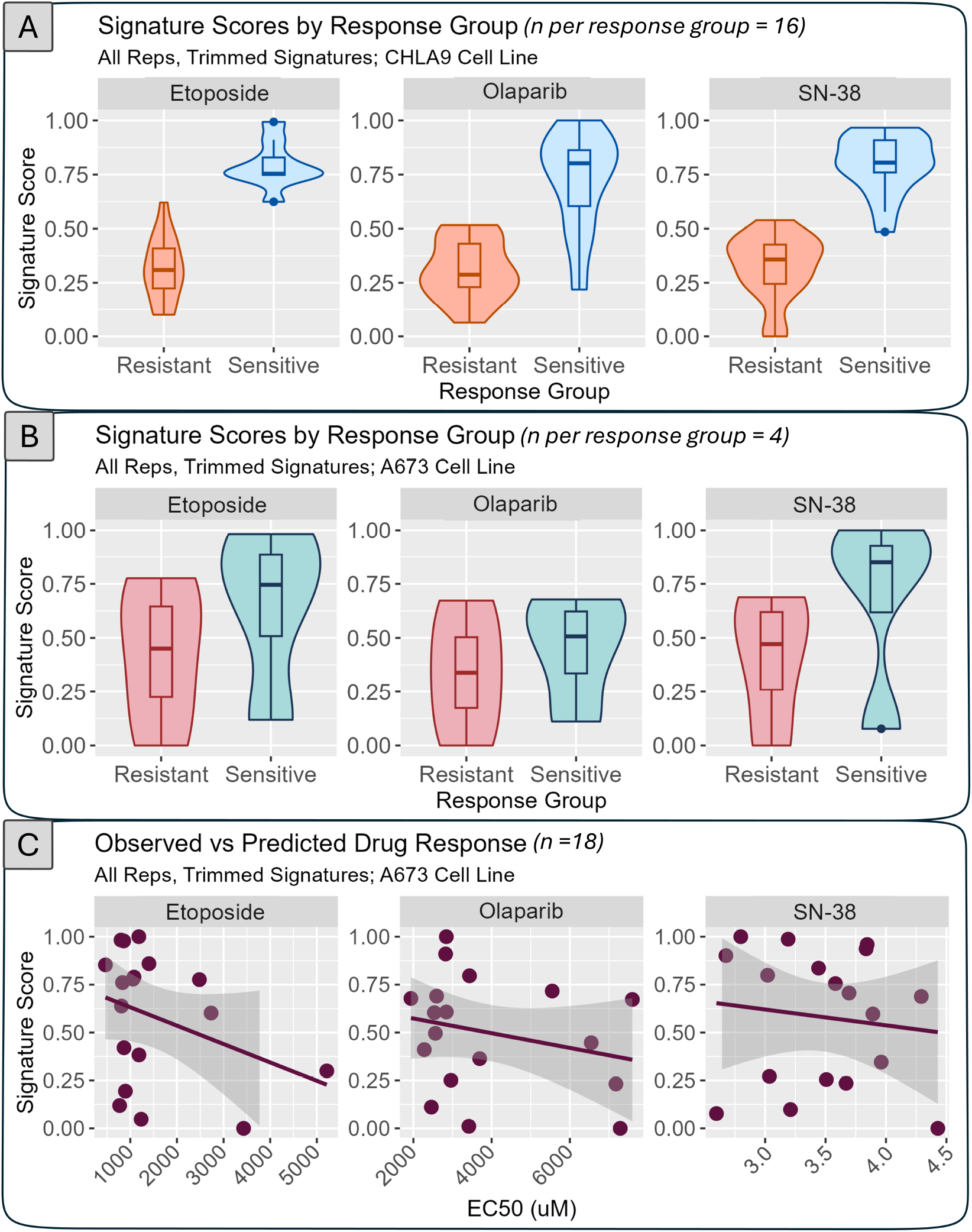
Signature scores between the most sensitive and most resistant samples. Signature scores (the median expression *z*-score of the signature genes, normalized across all cell lines within each dataset) for the 20% most resistant and sensitive samples are compared in different EWS cell lines. **Panel A** shows these values in CHLA9 samples, and **panel B** shows these results in A673 samples. The differences between resistant and sensitive groups are significant for all three signatures in panel A, but none are significant for panel B. **Panel C** shows signature scores for all A673 samples compared to their *EC*_50_s.

The signatures may not have been able to discretize sensitive and resistant samples as clearly in A673 compared to CHLA9 for three reasons. First, while A673 evolved resistance against the same drugs as CHLA9 in our study, the selection pressure differed somewhat. The doses of each drug in the combination and the concentration ratio between drugs within a combination differed greatly from the selection pressure we chose in this study, even though the drugs and alternating cycles remained the same between the two experiments. The doses also escalated as the pilot experiment proceeded, while the dose remained constant in this study. Second, the three signatures used in this comparison did not show the most promising results in CHLA9 relative to the other signatures, based on **Fig. 6B**. The etoposide signature had the least promising performance within the CHLA9 dataset (**Fig. 6B**). The performance of the SN-38 and olaparib signatures were only marginally better than that of the etoposide signature, but overall were still quite lacking. Lastly, the small sample size in the A673 study limited the statistical power of this analysis. Despite these limitations, our ability to observe the trend we expected between sensitive and resistant signature scores is promising and leads us to believe that even some of the more unremarkable signatures could still prove to be valuable with further validation.

## Discussion

Despite incremental gains in the management of Ewings sarcoma (EWS), outcomes for patients with relapsed or metastatic disease remain unacceptably poor. Roughly a quarter of patients with primary disease and 70% with metastatic disease relapse, and 5-year survival after relapse is *<* 15%.^1,3,5–7^ First- and second-line regimens rely on overlapping classes of alkylating agents and topoisomerase inhibitors, providing ample opportunity for cross-resistance.^20^ Given that many cancer-related deaths are caused by the evolution of drug resistance, there is a dire need for treatment strategies that explicitly anticipate and exploit tumor evolution.^31^ Gene expression signatures offer a straightforward, clinically translatable way to personalize therapy, as they can be used to predict drug response among different agents with a single biopsy. We can design signatures to forecast drug sensitivity in an evolution-informed manner by focusing on collateral response, where resistance to one agent coincides with increased sensitivity to another, and convergent evolution, where different organisms evolve similar phenotypes. The concept of convergent evolution can also be observed in cancer–even when tumors follow diverse evolutionary trajectories, they can arrive at phenotypically similar, drug-sensitive states.^16^ By identifying these convergent states and extracting shared transcriptional patterns among samples exhibiting collateral sensitivity, we increase the likelihood that our resulting biomarkers are robust to underlying heterogeneity.

To address the need for personalized therapy in EWS and generate robust predictive signatures, we first used experimental evolution to extensively capture evolutionary trajectories of EWS as drug resistance evolved. By evolving resistance repeatedly in an EWS cell line, CHLA9, under clinically relevant VDC-EC selection, measuring collateral drug responses, and quantifying gene expression over time, we generated a large dataset to identify rare but actionable phenotypes in a disease where patient-level resistance data are scarce. From this resource, we extracted predictive biomarkers of collateral drug response. Our signature extraction pipeline included 5-fold repeated subsampling, differential expression analysis, and stringent statistical trimming. Ultimately, our pipeline yielded 28 predictive collateral sensitivity signatures that outperformed random-gene null distributions in most tested models. Not all drugs produced equally informative signatures. For example, etoposide, a component of first-line therapy, showed little diversity in response, limiting our ability to detect sensitivity-associated transcripts. These observations suggest that a signature’s utility may depend on the breadth of collateral drug responses available for a given agent.

The extensive amount of data produced here holds potential utility outside the scope of this study, and all collected data have been made publicly available. This data could allow for closer inspection of the different mechanisms by which the evolution of resistance to VDC-EC may be occurring. Other mechanism-focused analyses could allow for further understanding of why collateral sensitivity is repeatable for some drugs but not others, and what drives the repeatability of evolution in this disease.

One limitation of our study was that our primary dataset is derived from *in vitro* evolution in a single cell line background, which cannot fully recapitulate the complexity of patient tumors. However, given that one of our primary aims was to maximize the number of potential evolutionary trajectories possible from a single origin, we decided to limit our study to a single cell line that was the most genomically similar to patients, compared to other EWS cell lines, and had not been exposed to treatment yet.

There are a few ways we may improve our predictive signature library, particularly to maximize clinical relevance. Our preliminary test in an independent EWS line (A673) reproduced the expected directionality of signature scores, but small sample size and modest performance for certain signatures highlight the need for larger, physiologically relevant datasets. Ideally, we would use patient-derived xenografts or *ex vivo* cultures paired with drug testing to rigorously assess the robustness of the signatures. Future refinement could incorporate clinical transcriptomes directly into the extraction process. For instance, our current gene sets could be filtered through co-expression networks derived from patient samples–an approach analogous to that of Buffa *et al* and Scarborough *et al* in their biomarker extraction methods.^16,32^ Retaining only highly co-expressed genes may enhance robustness and translatability.

Eventually, we envision a clinical workflow where a VDC-IE-resistant EWS tumor is biopsied, gene expression of the signature genes is quantified, and our library of collateral-sensitivity signatures is applied to generate a list of promising candidate drugs ranked by relative sensitivity, as we previously proposed.^33^ Such a framework could fit naturally into a broader framework of evolution-informed therapy, complementing other strategies aimed at preventing, delaying, or steering resistance.^34^

This work demonstrates that experimentally defining and exploiting convergent collateral sensitivity states can yield predictive gene signatures for a rare, phenotypically heterogeneous cancer currently lacking effective second-line treatment options. The data resource we generated, the analytic framework we developed, and the preliminary validation we performed collectively chart a path toward actionable, evolution-informed personalization of chemotherapy in EWS.

## Materials & Methods

### Cell Culture

CHLA9 cells (passage 20) were obtained from the Children’s Oncology Group’s biorepository. This cell line was derived from a 14-year-old female caucasian with a primary tumor in her thorax. The tumor had an EWS/FLI1 fusion mutation and a functional p53 gene, representative of the majority of EWS patients.^2^ CHLA9 was established from the treatment-naive tumor of this patient. Cell lines tested negative for mycoplasma before, at the midpoint, and after the experiment. Cells were cultured in Iscove’s Modified Dulbecco’s Medium (Cleveland Clinic Research Media Core) supplemented with 1% penicillin/streptomycin (Cleveland Clinic Research Media Core) and 20% fetal bovine serum (Gibco, A52567-01) within cell-culture-treated T75 flasks.

#### Evolving Resistance in EWS Cell Lines

Each treatment cycle began with a proliferation stage, during which the cells were allowed to grow without drug until 80-90% confluency in a T75 flask. After this point, we added either the vincristine-doxorubicin-cyclophosphamide (VDC) or the etoposide-cyclophosphamide (EC) combination, and the cells were exposed for five days, including the initial date of exposure. After five days, the cells were rinsed with phosphate-buffered saline, and fresh drug-free medium was replaced. Once the cells reached 80-90% confluency again, the replicate was divided into four groups of various cell densities depending on the task: one group was frozen at − 80°C, one group was seeded into a new T75 flask for the next cycle, one was set aside for RNA and DNA extraction, and the last group was seeded into two 384-well plates for collateral drug screening and VDC or EC drug response screening (**Fig.1B**).

Experimental evolutionary replicates were treated with the *EC*_70_ of wild-type CHLA9 against VDC (vincristine, doxorubicin, and cyclophosphamide) and EC (etoposide and cyclophosphamide). The VDC concentrations included 0.0462nM vincristine, 0.0277 uM doxorubicin, and 0.8418 uM cyclophosphamide. The EC concentrations included 0.184 uM etoposide and 3.945 uM cyclophosphamide. The ratio between each drug in the combination was determined by comparing the maximum observed plasma concentration of each individual drug.^18^ Vincristine, doxorubicin, and etoposide were dissolved and diluted in dimethyl sulfoxide (DMSO), and working stocks were created in advance of the experiment and stored at − 20°C until use. Cyclophosphamide (stored in 1 mg powder aliquots at −80°C) was reconstituted, activated, and diluted on an as-needed basis in a 1 mg/mL sodium thiosulfate solution in sterile water.

### VDC-EC Drug Response Screening & Analysis

Vincristine (item no. 11764), doxorubicin (item no. 15007), etoposide (item no. 12092), and 4-hydroxyperoxy-cyclophosphamide (item no. 19527, referred to in this paper as cyclophosphamide) were obtained from Cayman Chemicals. Working stocks for these drugs were created and aliquoted at the same time before the beginning of the experiment, and the same batch of stocks was used throughout the experiment. VDC drug response was measured at the end of every VDC cycle (odd-numbered cycles), and EC drug response was measured at the end of every EC cycle (even-numbered cycles). A 3-fold dilution with 12 concentrations (including no drug/vehicle only) was done in triplicate per evolutionary replicate in a cell-culture-treated 384-well plate (catalog no. 142761) at 5,000 cells per well. For VDC, the maximum concentrations were 3.742 nM vincristine, 2.244 uM doxorubicin, and 68.184 uM cyclophosphamide. For EC, the maximum concentrations were 44.631 uM etoposide and 959.649 uM cyclophosphamide. A solution of the maximum concentration was created and then diluted afterward so that the ratio between drugs remained constant across the serial dilution. The drug was serially diluted, then added the day after cells were seeded. Cells were exposed to the drugs for five days (including the initial day of exposure). After this time, (1/3)X Cell Titer Glo (ProMega, G9243) was added in equal volume per well, incubated for 10 minutes, and then luminescence was read using a TECAN Spark plate reader. The resulting data were normalized to percent survival using the no-drug control for each replicate. A 4-parameter log-logistic (Hill) function was then fit to each drug response curve in R (with a user-defined function shown in the GitHub repository). From this, the *EC*_50_ and *IC*_50_ were calculated.

### Collateral Drug Response Screening & Analysis

The 48 drugs used for the collateral screen were obtained from MedChemExpress and used by the Cleveland Clinic drug screening core. Catalog numbers are shown in **Supplemental File 1**. All collateral drug response plates (cell-culture treated 384-well plates, Corning, 07-000-074) were stamped by the drug screening core prior to the start of the experiment with a Hamilton Liquid Handler. All plates were heat-sealed individually, vacuum-sealed in groups of ten for each time point, and stored at − 20°C. To ensure storage at this temperature did not affect drug efficacy, the drug response of wild-type CHLA9 was measured with and without one additional freeze-thaw before beginning the experiment, and no differences were observed. One collateral response plate was used per evolutionary replicate at each time point, and all drugs and concentrations were tested in triplicate. On all plates, 10 mM staurosporin was used as a positive control and 0.3% DMSO was used as a negative control. Prior to use, plates were thawed at 37°C for 10-20 minutes, and cells were seeded using a Multidrop Reagent Dispenser. Cells were exposed to the drugs for five days (including the date of initial exposure), after which (1/3)X Cell Titer Glo was added in equal volume per well, incubated for ten minutes, then luminescence was read using a TECAN Spark plate reader.

On each plate, two concentrations were tested in triplicate, with one concentration being the wild-type *EC*_20_ against a drug, and the other concentration being the wild-type *EC*_80_. These concentrations were determined by measuring a 12-point dose-response (not including 0 uM) with a 10-fold serial dilution starting from 10 uM against each of the 48 drugs against wild-type CHLA9. Thirteen drugs were unresponsive to all concentrations tested, so 10 uM was used in place of *EC*_20_ and *EC*_80_ for these agents. To quantify the collateral response of a sample compared to wild-type later in the analysis, log_2_(fold-change) in area-under-the-curve (AUC) was calculated for the 35 drugs where two different concentrations were measured. For the 13 drugs only measured at 10 uM, log_2_(fold-change) in normalized luminescence was calculated instead of AUC.

### RNA Sequencing & Analysis

Bulk RNAseq was performed by the Cleveland Clinic Genomics Core with a sequencing depth of 20M reads per sample. Out of 101 samples generated from the evolution experiment, 73 were sequenced. 28 samples were not sequenced due to poor RNA quality and/or insufficient RNA quantity. Raw data was pre-processed using STAR on Case Western Reserve University’s High-Performance Computing Cluster.^35^ Low read numbers (≤ 50 reads) were filtered out from the count data and quantile-normalized with the EBSeq package in R.^24^ Differential expression was calculated using EBSeq.^24^

#### Signature Trimming Parameters

First, gene expression between top and bottom 20% of responders was compared for each individual gene using a *t-test*, and only genes yielding a *p-value* less than a Bonferroni-corrected *α* of 0.05 (corrected using the number of genes in the signature) were retained. With the remaining genes, we calculated Cliff’s Delta, a non-parametric measure of effect size, between the same sample groups using a threshold of |*δ* | > 0.8.^25^ The standard threshold for Cliff’s delta is typically |*δ* | > 0.47, so the threshold we used here was far more conservative.^25^ Finally, we fitted an elastic-net regularized logistic regression, optimizing the hyperparameters in a nested scheme.^26,27^ Outer folds were created via five-fold repeated subsampling to split the pool of resistant and sensitive samples in the same manner as described previously. An inner fold was created for each of the five folds, using leave-one-out cross-validation to train the model on 70% of the data and test it on the remaining 30%. The training split explored pairs of *α* and *λ*, and the pair yielding the highest AUC after testing was selected. To finalize the signature, we applied the one-standard-error rule.^27^ Among all outer-fold models whose AUCs fell within one standard error of the best observed, we selected the model with the smallest number of non-zero coefficients. The *α* and *λ* pair used for this model was then used to refit a final model on the complete dataset. The final trimmed signature included only the genes with non-zero coefficients. Signature lengths before and after trimming are depicted in **Figs. S3** and **S4**.

## Data and Code Availability

All data cleaning, analysis, and plotting were performed using R (Version 4.5.0) with RStudio.^36^ The code to download, clean, and analyze the data associated with this work is available on GitHub at github.com/kxlinr/ews-signatures.

## Acknowledgements

We sincerely thank the Carson Sarcoma Foundation and Alan B. Slifka Foundation for funding this work.

## Author contributions statement

K.L.R. contributed to experimental design, wrote all associated code, analyzed data, and wrote the manuscript. E.C. assisted with experimental workload. J.G.S. contributed to experimental design, analyzed data, and wrote the manuscript. All authors read and approved the manuscript.

## Supplemental Figures

**Figure S1.**
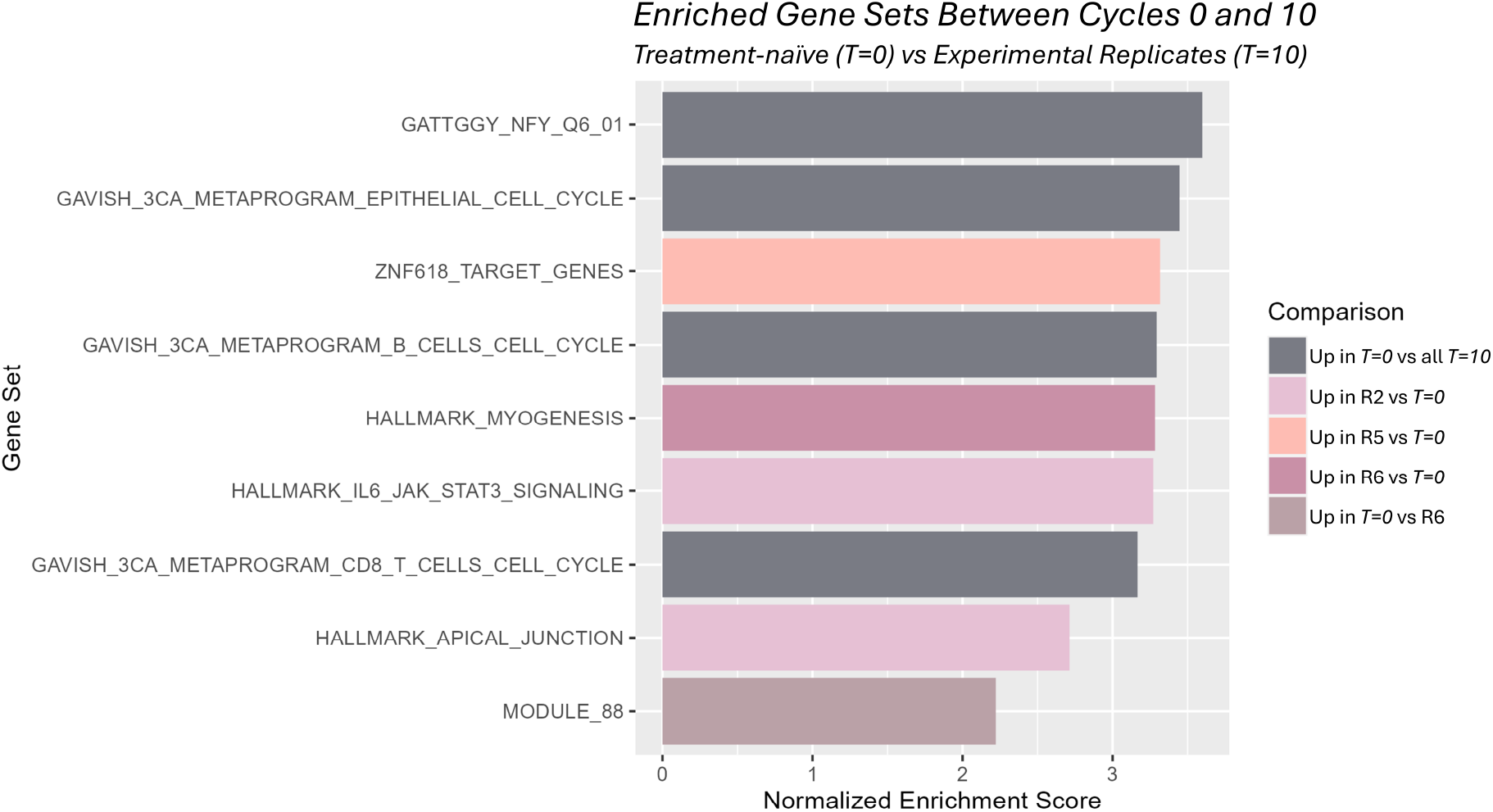
Gene set enrichment analysis between treatment-naive and treatment-resistant experimental replicates. Gene set enrichment was performed on differentially expressed genes between the treatment-naive cell line compared to experimental replicates at the end of the experiment that were sequenced (evolutionary replicates 2, 5, and 6), as well as the treatment-naive cell line compared to each of these replicates individually.

**Figure S2.**
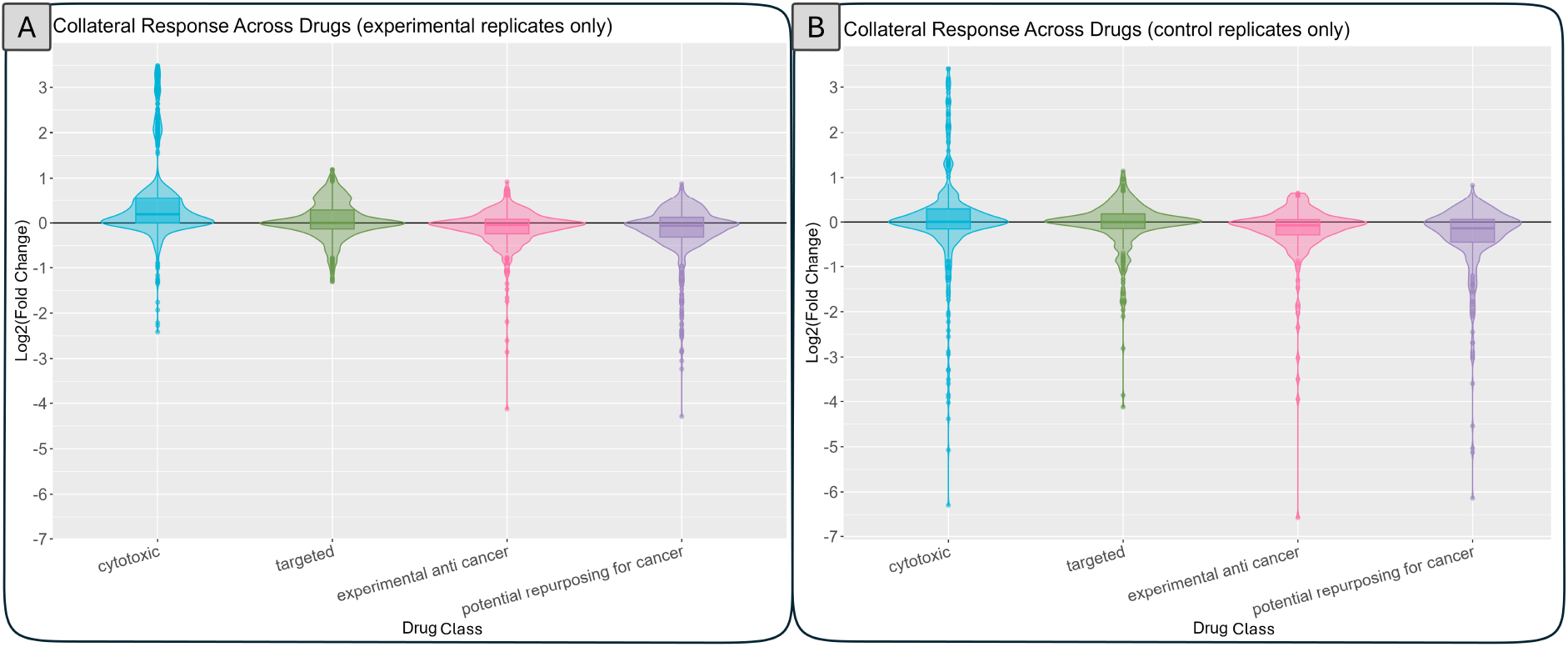
Collateral response by drug class. Collateral response to all samples is shown stratified by drug category. Each point within a distribution represents the log 2-fold change in response comparing wild-type to one evolutionary replicate at one point in time. Panel **A** shows experimental evolutionary replicates only, while panel **B** shows data in control replicates only. While differences between the groups within each panel are subtle, cytotoxic agents averaged the highest collateral resistance against these drugs. Conversely, experimental cancer drugs and other existing agents that are being investigated for repurposing against cancer have the most collateral sensitivity against those agents. Overall, the control replicates tended to be somewhat more sensitive against all agents compared to the experimental replicates, but largely showed similar distributions in drug response.

**Figure S3.**
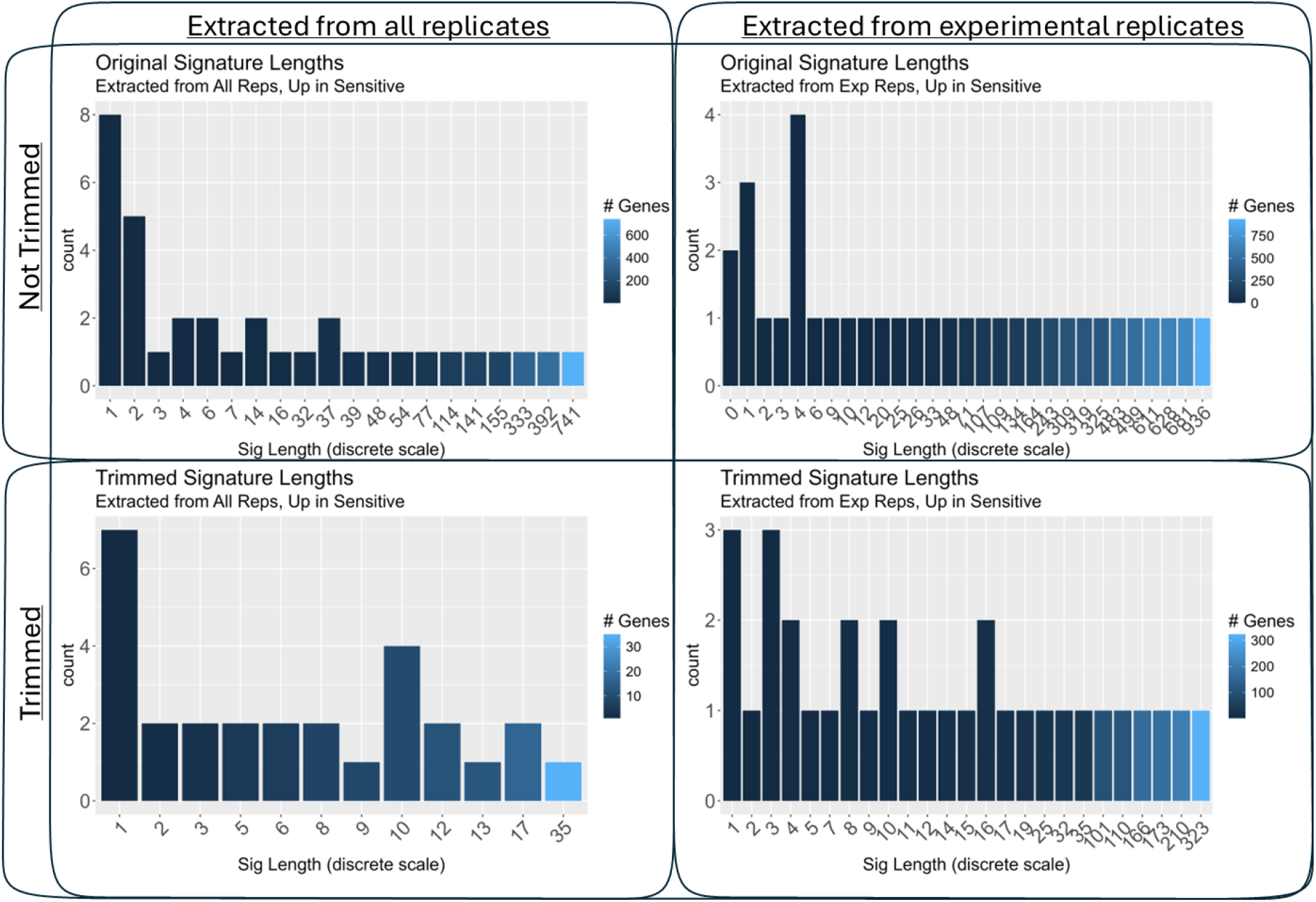
Distribution of up-regulated signature length before and after trimming. For all panels, the number of genes in a signature (with only genes that are up-regulated when a sample is sensitive) is shown on the x-axis as a discrete scale to maximize readability. The number of signatures with a certain number of genes is shown on the y-axis. Plots on the top row show these distributions without trimming the longer signatures, while plots on the bottom row show the distributions after trimming signatures over 50 genes. The two plots on the left depict signature lengths from the signatures extracted using all evolutionary replicates, including controls. The two plots on the right depict signature lengths from the signatures extracted using only experimental evolutionary replicates.

**Figure S4.**
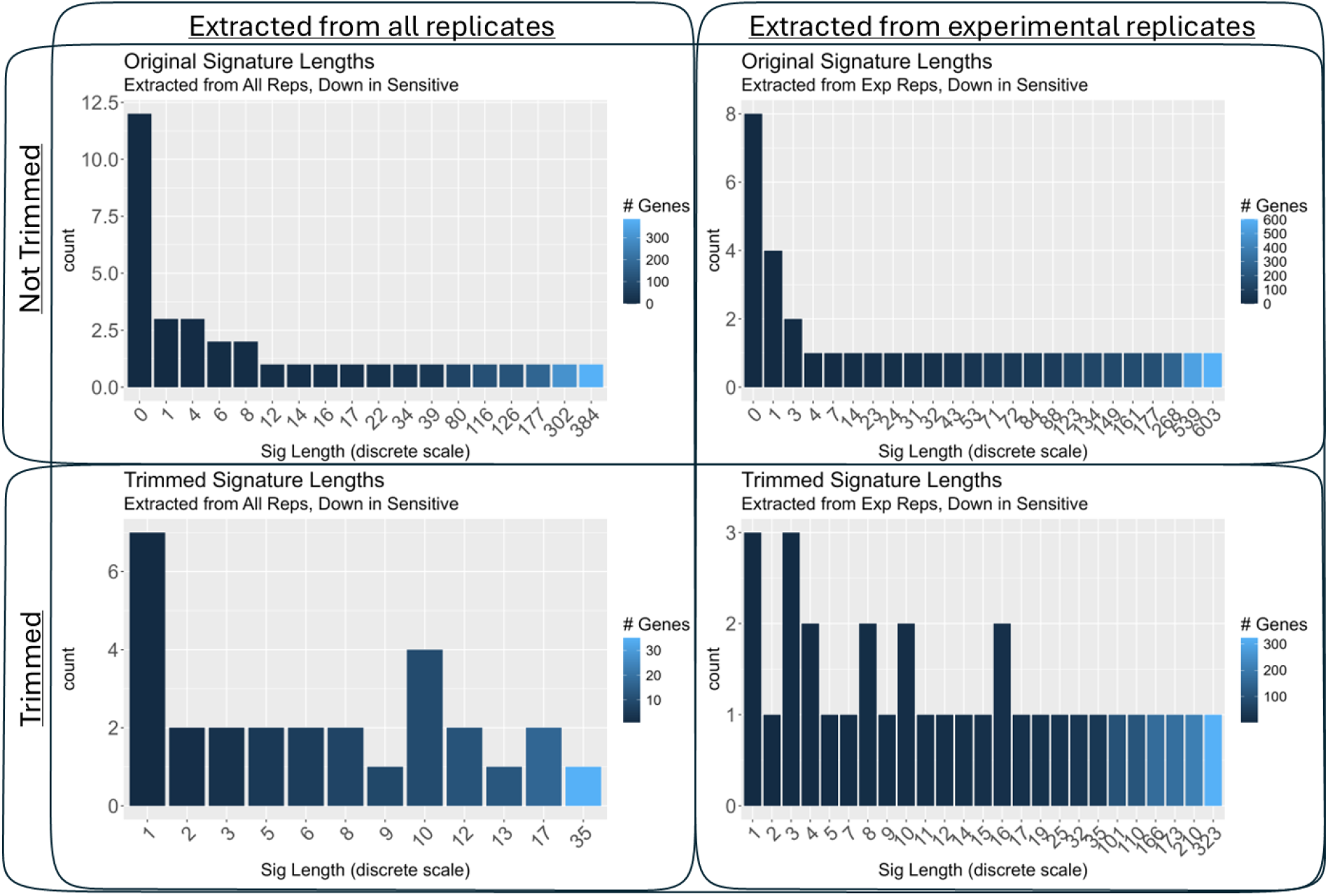
Distribution of down-regulated signature length before and after trimming. For all panels, the number of genes in a signature (with only genes that are down-regulated when a sample is sensitive) is shown on the x-axis as a discrete scale to maximize readability. The sub-panels are laid out in the same fashion as Fig. S3.

**Figure S5.**
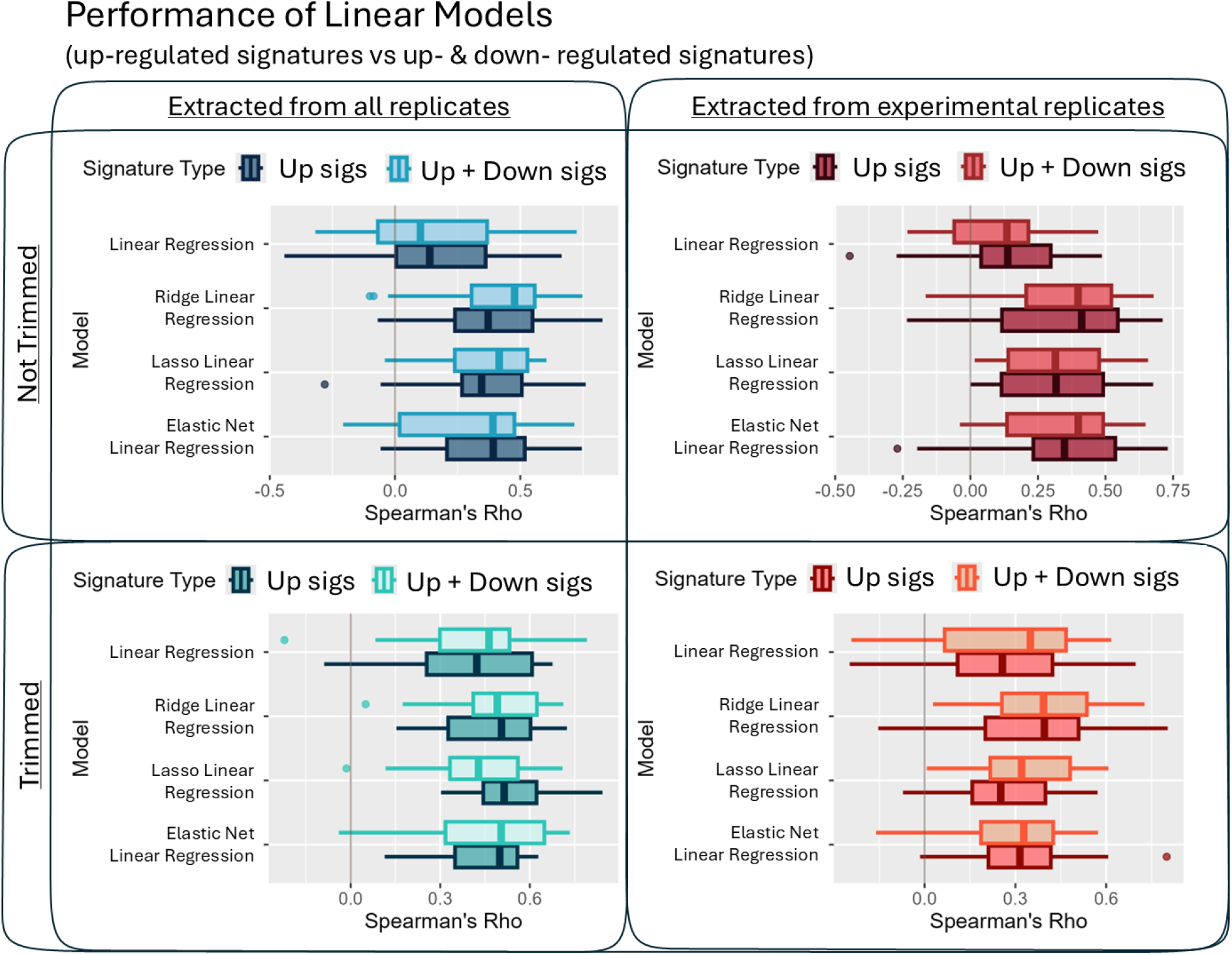
Performance of linear models trained on up-regulated gene signatures compared to up- and down-regulated signatures together. Spearman’s rho (shown along the x-axis) between predicted and observed survival is shown for four different linear models (linear regression, linear ridge regularization, linear lasso regularization, and linear elastic net regularization). Within each quadrant, the lighter colored boxplots show Spearman’s rho from models fitted with genes that are up-regulated or down-regulated in sensitive samples. The darker colored boxplots show Spearman’s rho from models fitted with only genes that are up-regulated in sensitive samples. Each quadrant shows signatures extracted from either all replicates or experimental replicates only, and with or without trimming. No statistically significant differences are found between up-regulated only and up- and down-regulated gene signatures in any case.

**Figure S6.**
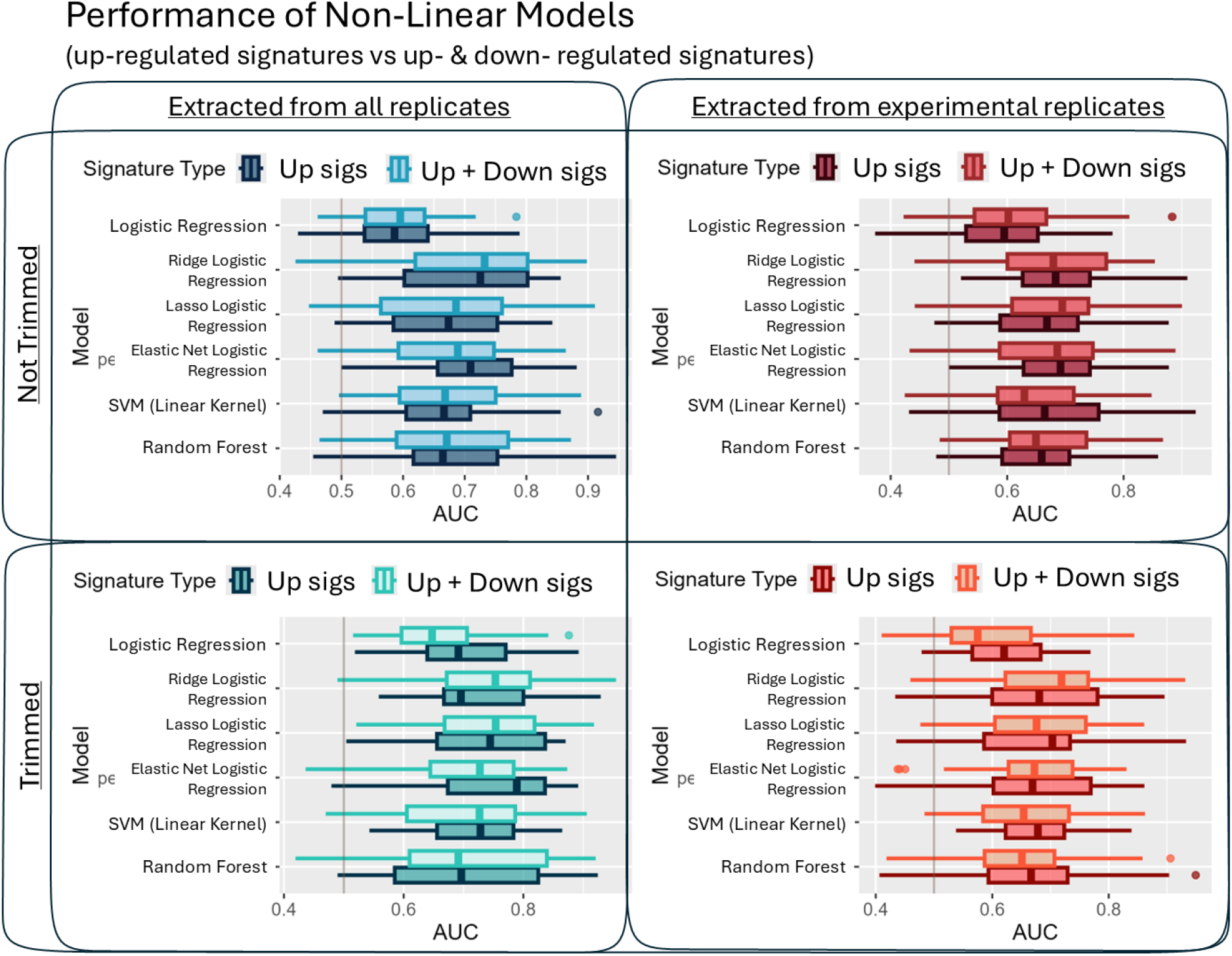
Performance of non-linear models trained on up-regulated gene signatures compared to up- and down-regulated signatures together. AUC (shown along the x-axis) between predicted and observed survival group is shown for six different non-linear models (logistic regression, logistic ridge regularization, logistic lasso regularization, logistic elastic net regularization, support vector machine with a linear kernel, and random forest classification). Within each quadrant, the lighter-colored boxplots show AUC from models fitted with genes that are up-regulated or down-regulated in sensitive samples. The darker-colored boxplots show AUC from models fitted with only genes that are up-regulated in sensitive samples. Each quadrant shows signatures extracted from either all replicates or experimental replicates only, and with or without trimming. No statistically significant differences are found between up-regulated only and up- and down-regulated gene signatures in any case.

**Figure S7.**
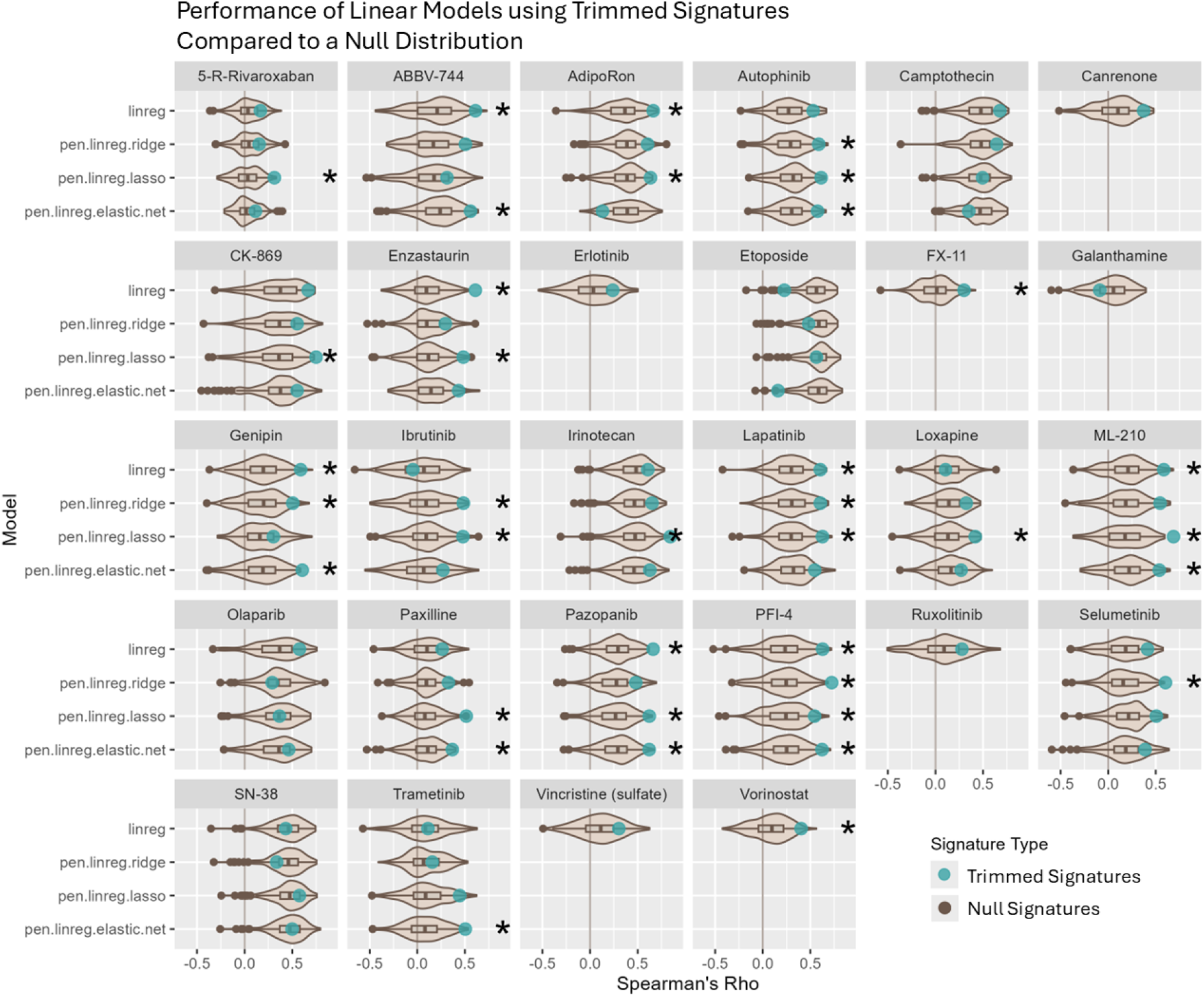
Performance of linear models trained on up-regulated trimmed gene signatures (extracted from all replicates) compared to null distributions. Spearman’s rho is indicated along the x-axis, and linear model types are indicated on the y-axis. Spearman’s rho for 200 null signatures (randomly generated and the same length as the actual signature) is shown in the brown boxplot-violin. Spearman’s rho for the actual signature is indicated by the turquoise point. Asterisks indicate a statistically significant difference between the signature’s rho and the null distribution based on a permutation test with *α* ≤ 0.05. Some signatures are missing null distributions and actual signature metrics because the models could not be fitted with the given signatures. Penalized regressions tend to fail when predictors are too similar to each other, and this was the case with several of our shorter signatures.

**Figure S8.**
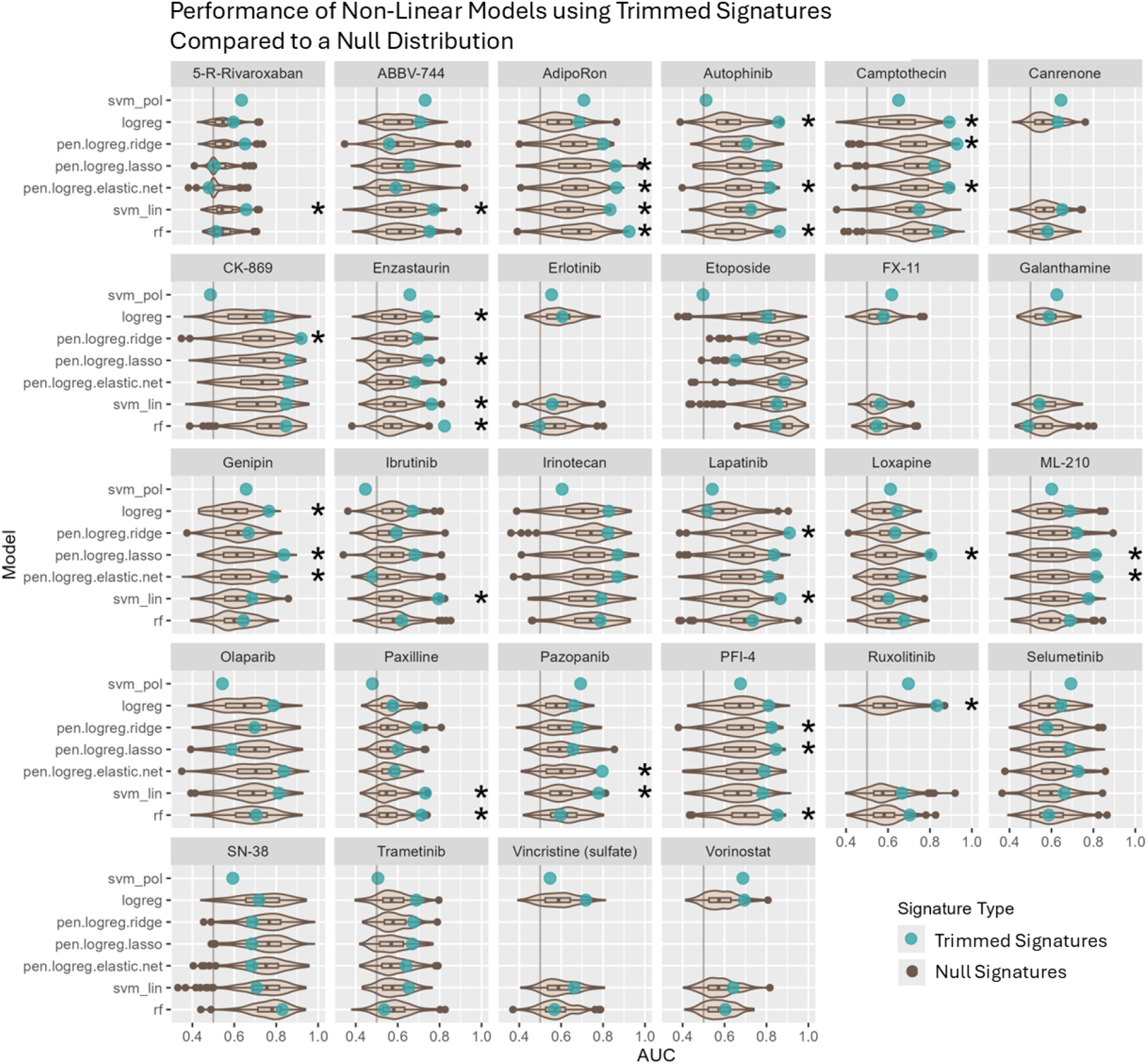
Performance of non-linear models trained on up-regulated trimmed gene signatures (extracted from all replicates) compared to null distributions. AUC is indicated along the x-axis, and linear model types are indicated on the y-axis. AUC for 200 null signatures (randomly generated and the same length as the actual signature) is shown in the brown boxplot-violin. AUC for the actual signature is indicated by the turquoise point. Asterisks indicate a statistically significant difference between the signature’s AUC and the null distribution based on a permutation test with *α* ≤ 0.05. Some signatures are missing null distributions and actual signature metrics because the models could not be fitted with the given signatures. Penalized regressions tend to fail when predictors are too similar to each other, and this was the case with several of our shorter signatures.

